# Learning efficient representations of environmental priors in working memory

**DOI:** 10.1101/2022.07.05.498889

**Authors:** Tahra L Eissa, Zachary P Kilpatrick

## Abstract

Experience shapes our expectations and helps us learn the structure of the environment. Inference models render such learning as a gradual refinement of the observer’s estimate of the environmental prior. For instance, when retaining an estimate of an object’s features in working memory, learned priors may bias the estimate in the direction of common feature values. Humans display such biases when retaining color estimates on short time intervals. We propose that these systematic biases emerge from modulation of synaptic connectivity in a neural circuit based on the experienced stimulus history, shaping the persistent and collective neural activity that encodes the stimulus estimate. Resulting neural activity attractors are aligned to common stimulus values. Using recently published human response data from a delayed-estimation task in which stimuli (colors) were drawn from a heterogeneous distribution that did not necessarily correspond with reported population biases, we confirm that most subjects’ response distributions are better described by experience-dependent learning models than by models with no learned biases. This work suggests that systematic limitations in working memory reflect efficient representations of inferred environmental structure, providing new insights into how humans integrate environmental knowledge into their cognitive strategies.

## Introduction

Traditional descriptions of working memory, a core feature of cognition [1], conceive of a system that takes in, maintains, and computes information over short timescales without a constant source of input. Knowing the limitations of this system can help identify its role in cognition [2] and provide a bridge to developing relevant neural theories. The limits and biases of working memory can be measured by the statistics of recall errors after a delay, for instance, in a visual delayed response task [3]. In these tasks, humans are asked to recall object features, such as location, color, or shape, a short time after presentation [2,4–6]. When feature values lie on a continuum, subject responses do as well, giving finely resolved measurements of the direction and magnitude of errors on each trial [7,8]. For example, people’s responses on delayed-response tasks often exhibit error magnitudes that increase roughly linearly with time, comparable to the variance of a diffusion process [9,10], providing a metric that can guide neural theories for working memory.

Complementary to behavioral studies of working memory, theories describing how the brain encodes information over short periods of time provide mechanistic insight. One well-validated theory associates remembered stimulus values with persistent neural activity in recurrently coupled excitatory neurons that are preferentially tuned to the target values [11]. Broadly tuned inhibitory neurons driven by excitation stabilize this activity into a localized structure called an activity *bump* [12,13]. Variability in neural tuning and synaptic connectivity can cause this activity bump to wander about feature space, causing trial-by-trial errors and biases often perceived as limitations to the system [14–16]. For example, delayed estimates may exhibit serial bias, whereby stimulus values from previous trials may attract or repel the retained memory of the most recent stimulus value [17]. Analogous attractive biases emerge when subjects retain the values of multiple stimuli within a single trial [18]. Additionally, subjects may exhibit systematic biases that include preferences for focal colors [16,19], orientations [20] and cardinal directions [21].

While biases are often considered reflections of suboptimality, they can be advantageous when reflecting the structure of the environment or sequences of stimuli the subject might see [22, 23]. There is ample evidence that the working memory system can be trained, and such biases may be the result of long-term learning [24]. Mechanistically, systematic biases in stimulus coding or delayed estimates could emerge from variation in the sensitivity of stimulus feature tuning across neurons [6,25,26]. Alternatively, the strength of synaptic connectivity may vary systematically so collective neural activity is biased to specific conformations in the network [27]. Such heterogeneity in the spatial organization of synaptic connectivity can reduce error by maintaining representations that are less susceptible to noise perturbations [28–30]. Thus, if synaptic heterogeneity reflects the expected distribution of stimulus values, recall of common features would be less error prone, improving cognitive efficiency [31].

Since certain stimulus features may be overrepresented in the natural world (e.g.., green/brown colors are more common in a forest; see also [32, 33]), we propose that subjects’ systematic biases could result from learning the natural distribution of specific features of the environment, which modulates synaptic connectivity to produce representation biases. Here, we model the effects of environmental feature distributions on delayed estimation in neural circuit models and their low-dimensional reductions, considering both models with network connectivity that is fixed and those shaped by long-term plasticity. We compare these results to human behavior and find that most subjects exhibit strategies best described by learning models, supporting the hypothesis that long-term representation biases are the result from learning environmental structure.

## Results

We begin with the premise that features of natural environments appear according to distributions that favor particular values that are overrepresented and thus, statistically more likely to occur (Fig. **1a**). Such parametric distributions could take on general forms [26], but for illustration, we assume a parametric prior that is periodic with peaks (dips) at common (rare) stimulus values

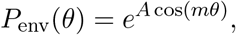

where *θ* corresponds with a particular feature value described on a ring (e.g., one feature dimension wraps periodically), *A* describes the amplitude, and *m* describes the number of peaks in the probability distribution. This periodic function resembles the color biases displayed by humans in [16] as well as cardinal bias common to angle and direction estimates [20,21]. Note, unless otherwise stated, all subsequent results assume that *m* = 4, so the peaks are centered at cardinal angles of *θ*, and *θ* is in radians in formulas, but plotted in degrees in figures for readability.

**Figure 1:**
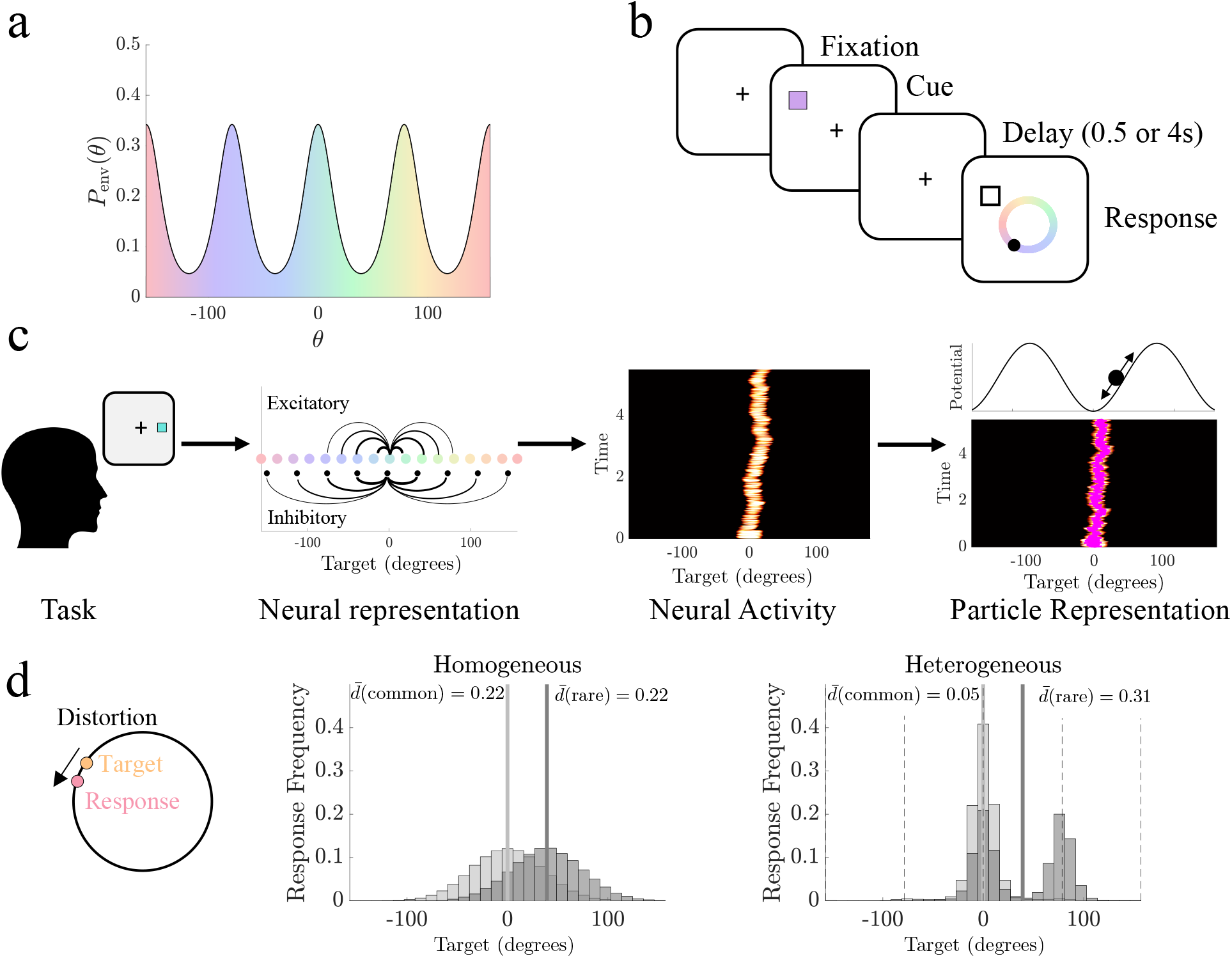
Heterogeneity in the distribution of environmental features is reflected in delayed estimates. **a.** Natural environments that include overrepresented features, such as certain colors, are described by heterogeneous priors of feature distributions with peaks at the overrepresented features. **b.** Schematic of the delayed estimation task, which requires subjects to remember a target feature (e.g., color) and report it following a short delay period. **c.** The remembered target feature is represented neuromechanistically by a subpopulation of stimulus-specific excitatory neurons with local recurrent excitatory and broad inhibition. The target representation is retained as a bump of sustained neural activity wandering stochastically during the delay. Bump dynamics can be projected to a particle model describing its stochastically evolving position: Spatial heterogeneity in synaptic connectivity is inherited by the particle model as a nontrivial energy landscape with attractors corresponding to regions of enhanced excitation. **d.** Distortion, the circular distance between the target and responses, is influenced by synaptic heterogeneity. In homogeneous networks, response errors at common environmental targets (*θ* = 0, light grey) and rare targets (*θ* = 45, dark grey) are equivalent, giving the same local mean distortion 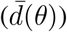. With synaptic heterogeneity matched to the environmental prior *P_env_*(*θ*), errors are reduced near common stimulus feature values (dashed lines). Parameters used as listed in Methods Table 1.

Our models describe the maintenance of estimates of continuous features [34,35], arising in tasks where an observer is briefly shown a number of items and, after a delay, probed about remembered stimulus feature values (e.g., location, orientation, or color). These models allow us to theorize how the true priors on the environmental features impact (and potentially bias) how stimulus feature values are remembered (Fig. **1b**). To illustrate, we focus on examples in which subjects recall colors, though equivalent results can be produced for models of orientation and location recall. Our models are motivated by previous observations that show human performance on delayed estimation tasks degrades over time, such that response variance increases roughly linearly, suggesting a diffusive process drives memory errors [9]. Such diffusive degradation of a stimulus estimate has been modeled in neural circuits as a localized region of persistent activity (bump) that stochastically wanders feature space due to neural and synaptic fluctuations [11,36]. Activity bumps emerge from strong stimulus-tuned recurrent excitation paired with broad stabilizing inhibition, which generates self-sustained activity [12]. Spatial variation in synaptic connectivity can shape the preferred locations (attractors) of the bump, introducing drift toward attractors.

Since the location of the activity bump is a proxy for the remembered stimulus feature value [14,15], we can simplify our analysis of the impact of activity bump fluctuations by considering low-dimensional models that describe the bump as a particle stochastically moving through an energy landscape (Fig. **1c**). Spatial variations in synaptic connectivity are thus accounted by the resulting energy landscape (and bump drift) they invoke [28,29,37] and can be derived asymptotically [37,38], making it a useful simplification of delayed estimate dynamics. Energy landscapes can be updated to represent an observer’s current estimate of the environmental feature distribution *P*_env_(*θ*) (see Methods) and can be more easily fit to response data than neural circuit models [14,16,23,39], providing a tractable model for studying the origins of systematic biases in working memory.

We compute our models’ average error between a true target feature value *θ* and its estimate as the mean distortion 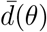, the circular distance between the target and responses. Overall error across all target values is computed as the total mean distortion 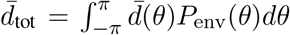 [15,29]. Thus, when synaptic connectivity (and the corresponding energy landscape) is aligned with the environmental prior *P*_env_(*θ*), the mean distortion is reduced at common target feature values 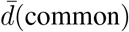 but increased for rare values 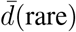. In contrast, purely distance-dependent synaptic connectivity (and a flat energy landscape) produces response distributions and mean distortion that are similar for common and rare target feature values (Fig. **1d**), making mean distortion a useful metric for quantifying error with respect to changes in synaptic connectivity.

Combining analysis of the energy landscapes with our distortion metric, we now systematically consider the impacts of environmental stimulus distributions on working memory responses, which can guide our understanding of how expectations about the environmental prior can be learned from experience and how these expectations can lead to more efficiently retained memories.

### Energy Landscapes Shape Recall Distortion

#### Uniform Stimulus Priors

We consider a particle model that describes the stochastically evolving estimate of the target feature value with an energy landscape that can incorporate bias, introduced by breaking the symmetry of continuous attractor models of delayed estimation [40]. This low-dimensional model can be derived asymptotically from the stochastic evolution of the position *θ*(*t*) of an activity bump that encodes the estimate and the information about the prior in its network connectivity (see Methods). The energy landscape that reflects an observer’s long-term estimate of the periodically-varying environmental prior *P*_env_(*θ*) can be generated as

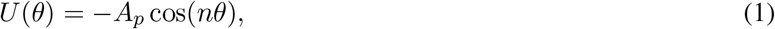

where *A_p_* describes the well amplitude and *n* is the number of attractors (each located at the believed common environmental feature values). This simple form for *U*(*θ*) allows us to probe how the alignment of the energy landscape to the true stimulus distribution shapes an observer’s distortion and produces response biases (Fig. **2a**).

**Figure 2:**
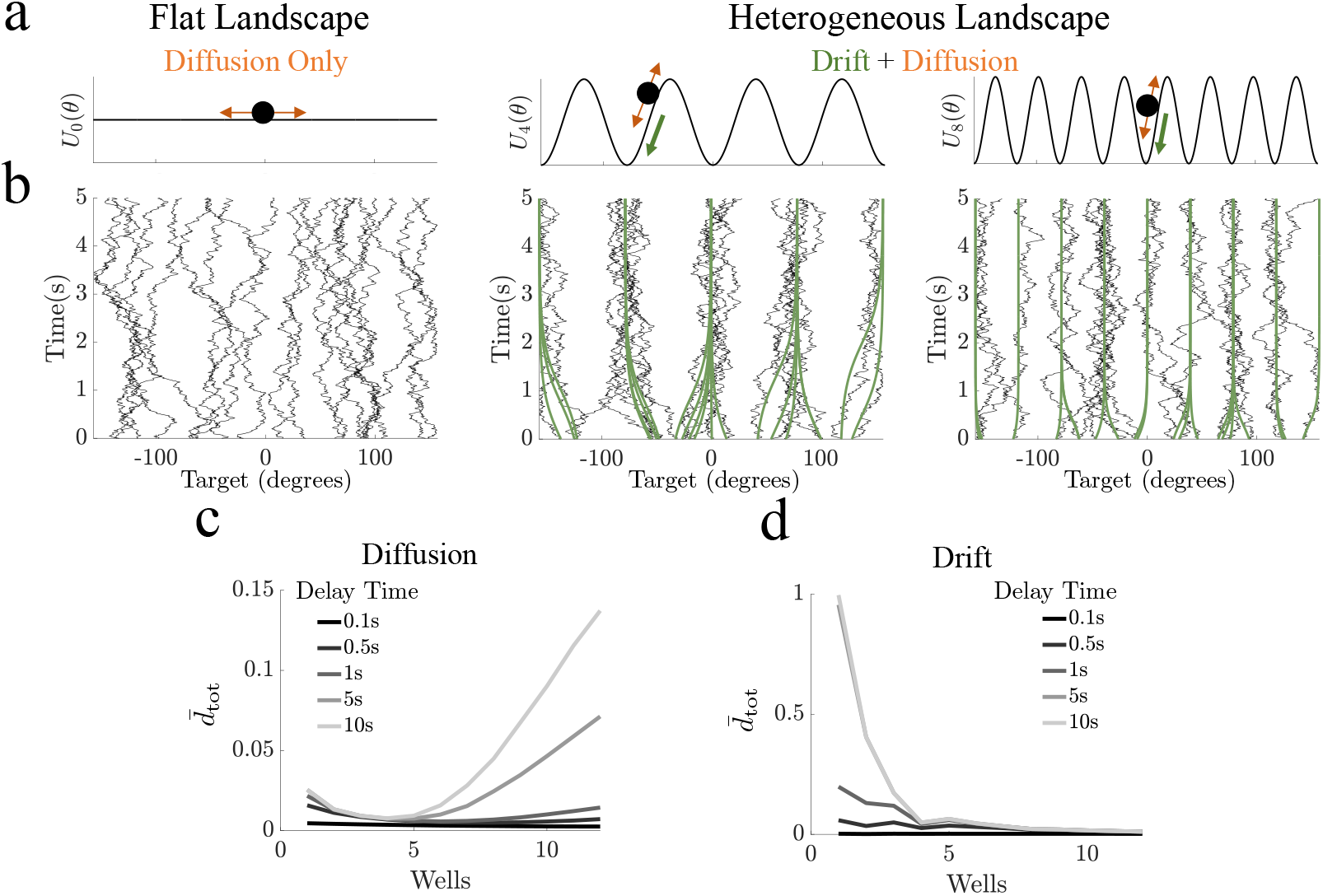
Particles in heterogeneous landscapes are drawn toward attractors. **a.** Schematics of the flat (homogeneous) landscape, with only diffusion, and heterogeneous landscapes, with potential-driven drift and diffusion. **b.** Example of particle trajectories in the flat and heterogeneous energy landscapes in **a**, sampled from a uniform environmental distribution. In the flat energy landscape, particle motion is driven purely by diffusion. In the heterogeneous energy landscapes, the particle drifts toward attractors over time but diffusion can cause the particle to “jump” wells. Drift-only process shown in green for comparison. **c.** Total mean distortion in a heterogeneous landscape with only diffusion (targets sampled at attractor points) and moderate diffusion (*σ* = 0.05). **d.** Total mean distortion in a heterogeneous landscape with only drift. Parameters for all sub-figures: *A_p_* = 0,1, *n* = 4,8, all others as listed in Methods Table 1.

The movement of the particle through this landscape evolves according to the stochastic differential equation

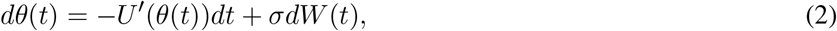

where the particle evolves as it: 1. descends wells surrounding attractors (drift), and 2. diffuses due to noise fluctuations given by *W*(*t*), a Wiener process. Considering a particle model with a flat energy landscapes (*A_p_* ≡ 0), memory of the target stimulus feature evolves according to pure diffusion during the delay period. In contrast, particles evolving along non-trivial energy landscapes (*A_p_* > 0) are biased toward the periodically placed attractors at *θ* = ±(*j/n*)*π* (*j* = 0,…,*n* – 1) (Fig. **2b**).

We first quantify the total mean distortion 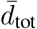 of responses from particle models encoding stimuli from a uniform prior. Even given a uniform environmental prior, delayed estimates can be improved due to the stabilizing effects of local attractors that mitigate the wandering from diffusion [28,29,41]. However, distortion of the target estimate is also enhanced by the introduction of drift. Therefore, we consider the individual contributions of diffusion and drift to the total mean distortion. Fixing the diffusion coefficient and sampling responses that only originate at attractor points (wells, e.g., *θ* = 0), we assess the impact of diffusion alone in heterogeneous energy landscapes (*n* > 0). Distortion varies non-monotonically with the number of wells (Fig. **2c**). This relationship is enhanced with longer delays and can be attributed to the close proximity of nearby wells as the well number (*n*) increases, reducing the strength of perturbation needed for the particles to ‘jump’ between attractors (see also Fig. **S1** and [29,42]). Low (high) levels of diffusion correspondingly show reduced (enhanced) total mean distortion (Fig. **S2**). In contrast, when we remove noise so particles in heterogeneous landscapes only drift, total mean distortion decreases considerably as the number of wells increases (Fig. **2d**), indicating reduced distortion is due primarily to the the constraint of diffusion by attractors.

#### Heterogeneous Stimulus Priors

In addition to the form of the energy landscape, mean distortion is impacted by the form of the environmental prior *P_env_* (*θ*). While the conditional probability of responses *P*(*θ*_resp_|*θ*_env_) is only altered by heterogeneity in the energy landscape, the marginal probability of response *P*(*θ*_resp_) is impacted by both the energy landscape and the environmental prior (Fig. **S3**), confirming that the mean distortion changes with the environmental prior. Matching the number and position of energy landscape wells to the peaks in the prior, we find the mean distortion 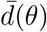 is significantly reduced at common (attractor) locations compared to a model with a flat energy landscape, but shows comparable levels of distortion at rare (saddle) locations (Fig. **3a**; bootstrapped distortion, *p* < 0.05).

**Figure 3:**
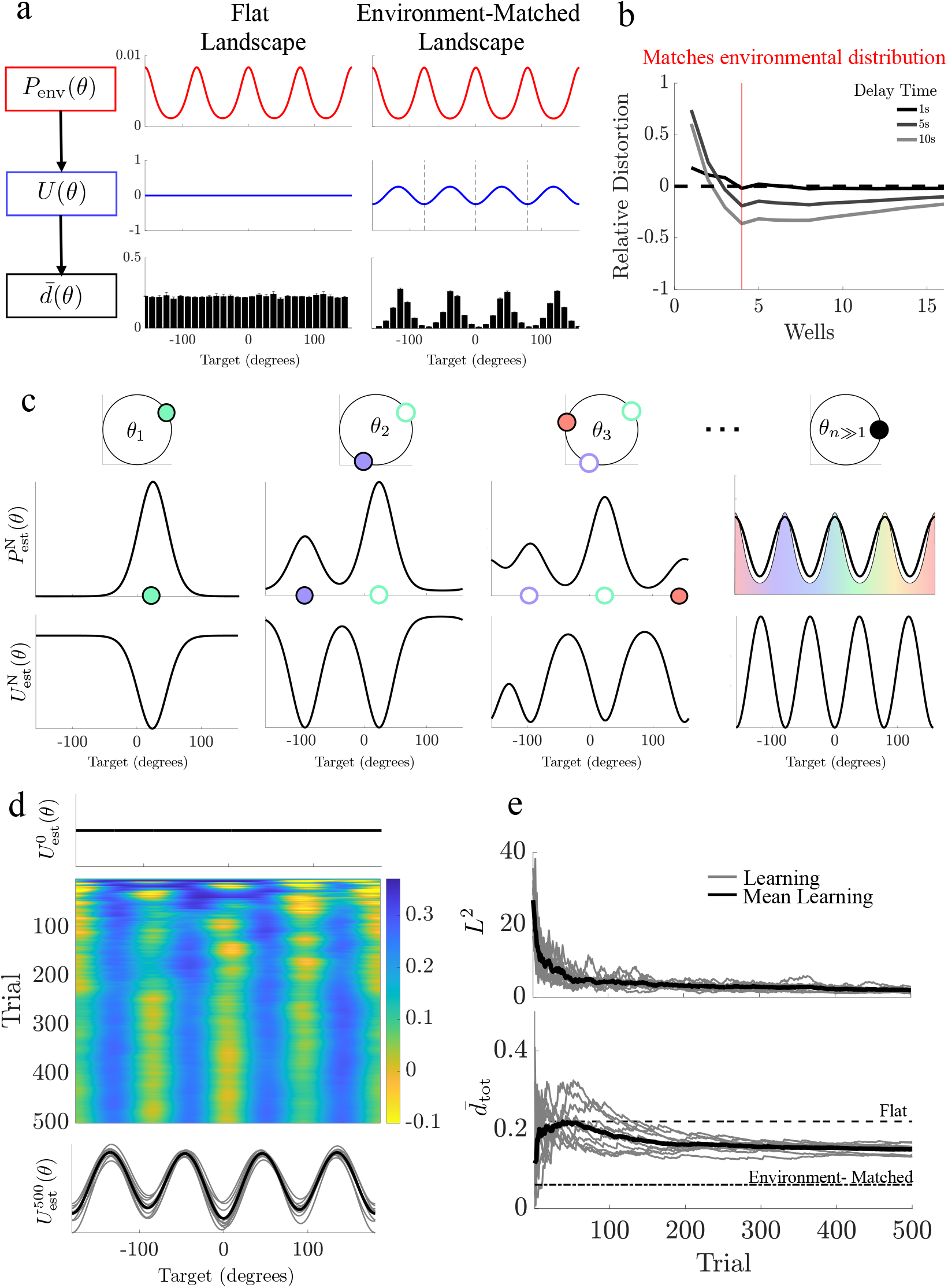
Distortion is reduced when energy landscapes match the environmental prior. **a.** Top: a heterogeneous environmental distribution (*P*_env_(*θ*)) passes through a heterogeneous energy landscape (*U*(*θ*)) and alters the mean bootstrapped distortion 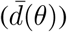 at a given target value *θ* (*N*_boot_ = 1*e*3). **b.** Relative total mean distortion compared between the flat landscape and heterogeneous landscapes (negative values denote reduced distortion for heterogeneous landscape). Red vertical line denotes where the number (and position) of attractors are aligned to peaks in the environmental prior. **c** Schematic of learning in a particle model. Based on the target observed on each trial, the estimated environmental distribution 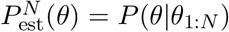 and the particle landscape 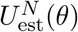 is updated at the target location. Over the course of many trials, the estimated distribution 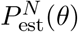 becomes more similar to the environmental prior *P*_env_ (*θ*), and the energy landscape aligns its wells to its peaks. **d** Heatmap showing the landscape updating over the course of many trials. Top trace shows initial landscape. Bottom trace displays the landscape on the final trial. Grey traces are 10 examples of the learning model, black trace is the average learning model’s landscape. **e** Top: *L*^2^–norm for the difference between the experience-dependent belief about the environmental distribution 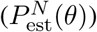 and the true environmental distribution (*P*_env_ (*θ*)). Bottom: Running average of the learning model’s total mean distortion 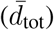. Parameters for all sub-figures as listed in Methods Table 1.

We ask if the total mean distortion typically decreases for periodic energy landscapes Eq. (1) as compared to flat landscapes when environmental priors are heterogeneous, and find that it is generally reduced (relative distortion is negative), with a dramatic reduction in distortion when attractors are aligned with peaks in the prior (Fig. **3b**), though the number of wells does not need to exactly match the number of peaks (Fig. **S4**). Energy landscapes misaligned with the environmental prior (e.g., aligned with rare target locations) generally produced response distributions with higher total mean distortion than aligned models (**Fig. S5)**, confirming that aligning attractors to environmental peaks increases coding accuracy of delayed estimates.

#### Experience-dependent Learning in Particle Models

We next ask whether energy landscapes that model the effects of long-term plasticity can infer a prior based on a long sequence of observations. The effective learning rule assumes subjects sequentially infer the environmental prior from long-term experience: After each trial, the subject’s running estimate of the environmental prior is merged with a likelihood function peaked at the current trial’s target value. This evolving estimate of the prior can be represented in the energy landscape by updating the landscape such that peaks in the prior estimate are encoded by attractors, corresponding to regions of synaptic potentiation in an equivalent neural circuit description (see Methods and Fig. **3c**). Over many trials, the energy landscape develops attractors aligned with the common feature locations (Fig. **3c,d**), regardless of observation order (Fig. **S6**). Thus, the experience-dependent updates generate learning of the environmental prior, and the energy landscape reflects better estimates of the environmental structure, which reduces total mean distortion, trending towards the distortion of a particle model assigned an environment-matched energy landscape (Fig. **3e**).

#### Subjects’ Behavior shows Hallmarks of Learning

We next validate our static and learning particle models against responses from a previously reported data set in which 120 human subjects perform sequences of delayed-estimation trials for target colors drawn from distributions along a one-dimensional ring (see [16] for more details). Subjects were cued with two items, the target and distractor, and asked to respond with the color of one item after a short (0.5s) or long (4s) delay. Item colors on each trial were selected from either an (a) uniform stimulus distribution or (b) heterogeneous distribution with four peaks, offset randomly for each subject (Fig. **4a**).

**Figure 4:**
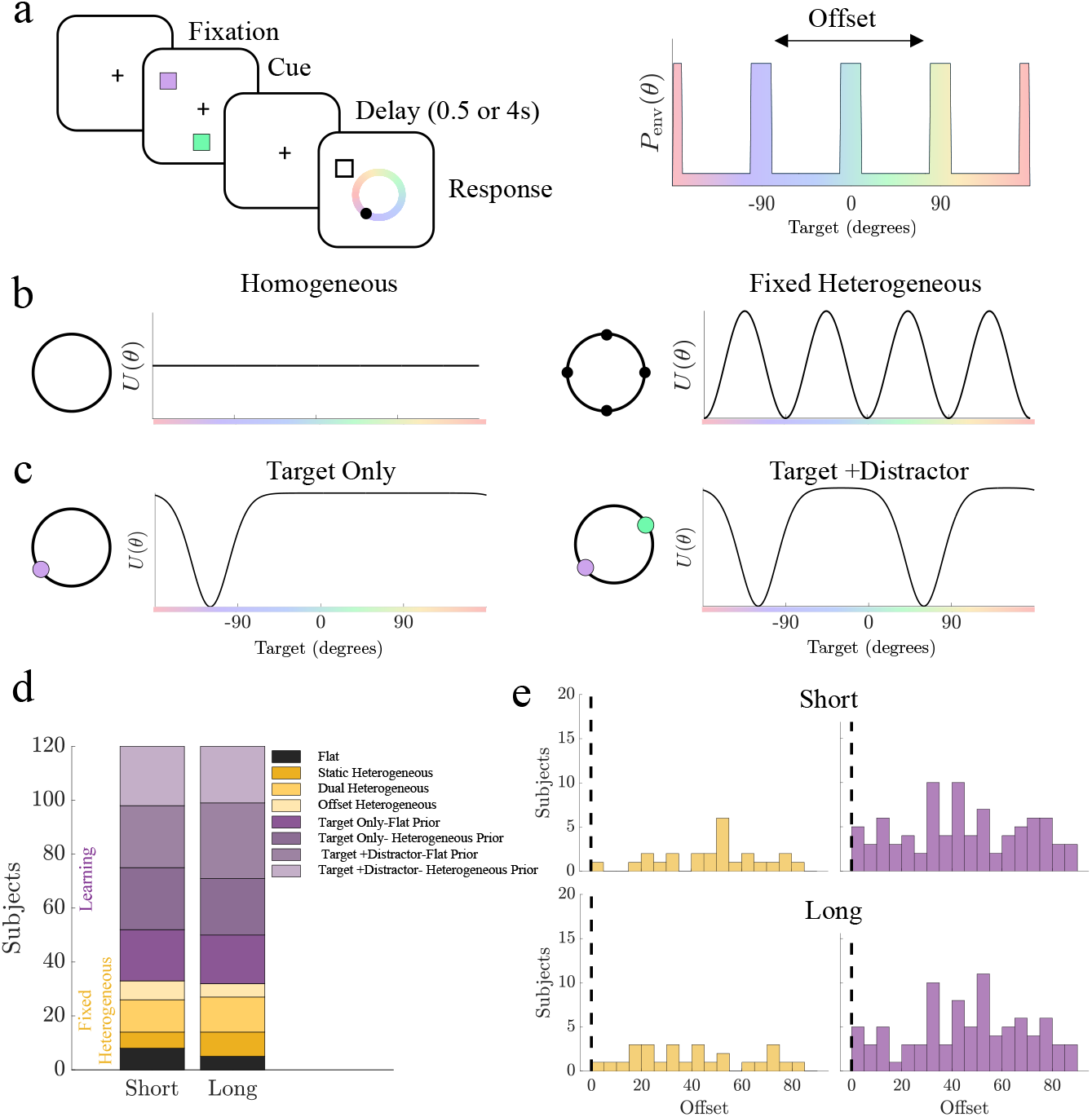
Subject responses based on targets drawn from heterogeneous distributions are best replicated by models governed by heterogeneous landscapes (static and learned). **a.** Experiment 2 from [16] in which subjects were shown two items, each of which could be drawn from a heterogeneous distribution whose peaks were evenly distributed but randomly offset for each subject. Subjects were prompted to respond with the corresponding color of one item. **b.** Fixed particle models used. Homogeneous landscape includes one free parameter, diffusion. Fixed Heterogeneous models include at least three free parameters: amplitude, number of wells, and diffusion. **c.** Learning particle models. Each model updates iteratively based on three parameters: width of the bump, depth of the bump, and diffusion. Target-Only learning models incorporate only the target prompted for response, and Target+ Distractor models incorporate both items. **d.** Number of subjects best matched to each model type for short and long delay periods. **e.** Assigned offsets for subjects best matched to Fixed Heterogeneous models (yellow) and Learning models (purple). Dashed lines shows human population bias location.

We ask if subject responses are best described by particle models with energy landscapes from one of three classes: (a) fixed and uniform; (b) fixed and heterogeneous; or (c) evolving from each subject’s stimulus history. Our fixed and heterogeneous class of models includes three variations: 1. a model with attractors spaced evenly around the ring aligned to each subject’s assigned environmental offset (Static Heterogeneous), with free parameters for the amplitude, the number of attractors, and the noise amplitude; 2. a variation allowing the offset of the attractors to deviate from the peaks of the prior (Offset Heterogeneous model); and 3. a variation in which the energy landscape is determined by two Fourier modes (Dual Heterogeneous model) (Figs. **4b** and **S7**).

We also consider four learning models: one form (two models) updates the energy landscape based only on the target (Target Only), and another form (two models) updates the potential landscape based on both observed items (Target + Distractor)(Fig. **4c**). The initial prior (initial landscape) is also varied to account for subjects’ potential systematic biases, since subjects can exhibit color biases even given uniform environmental priors [16]. Learning models are initialized either with a flat landscape (Flat Prior) or with a landscape with attractors at the locations of the subject population’s biases identified in [16] (Heterogeneous Prior) (Fig. **S7**).

To identify the model that best matches each subject’s responses, we apply cross-validation based on the mean squared error between subject and simulated responses (see Methods), across many possible parameter sets for each model. Nearly all subjects’ responses (93% of subjects in short trials and 96% of subjects in long trials) are best described by heterogeneous models, with a majority of subjects applying learning models (72% of subjects in short trials and 73% of subjects in long trials; Fig. **4d**). We find that many subjects best matched to learning and fixed heterogeneous models have assigned environmental prior offsets centered away from the population biases but not uniformly distributed for both short and long trials (p < 0.05, two-sample Kolmogorov-Smirnov test) and that the distribution of assigned offsets for learning model subjects is significantly different than that of subjects best fit to fixed heterogeneous models (p < 0.05, two-sample Kolmogorov-Smirnov test), with more learning model subjects having an assigned offset that is far from the population biases (Fig. **4e**). This finding suggests that many subjects confronted with observations from an environmental prior that differs from their baseline prior learn the new distribution of stimuli through experience.

#### Neural Mechanism for Learning Environmental Priors

We next identify a neural network model capable of implementing experience-dependent inference of environmental priors, comparable to our particle models [22,38] (see Methods for a demonstration that this model can be asymptotically reduced to our particle models). In this neural field model, excitatory neurons are tuned to preferentially activate given specific stimulus feature values. Average neural activity of neurons with a particular feature preference (location) *x* at time *t* is described by the variable *u*(*x,t*) with synaptic connectivity (combining excitation and inhibition) described by the function *w*(*x, y*). Excitatory coupling is strongest between neurons with similar preferential tuning, and inhibitory-excitatory feedback shows corresponding broader connections (Fig. **5a**; left). Combined local excitation and lateral inhibition supports the formation of persistent neural activity bumps when a transient input is presented at a particular location [11, 29, 43]. When synaptic connectivity depends only on the difference between neurons’ stimulus preferences, bumps have no intrinsically preferred positions in the network and lie along a continuous attractor, establishing an unbiased code for delayed estimation of an input stimulus value. Fluctuations that reflect synaptic noise cause bumps to wander freely with pure diffusion, so bumps are equally likely to move any direction (Fig. **5b**; left).

**Figure 5:**
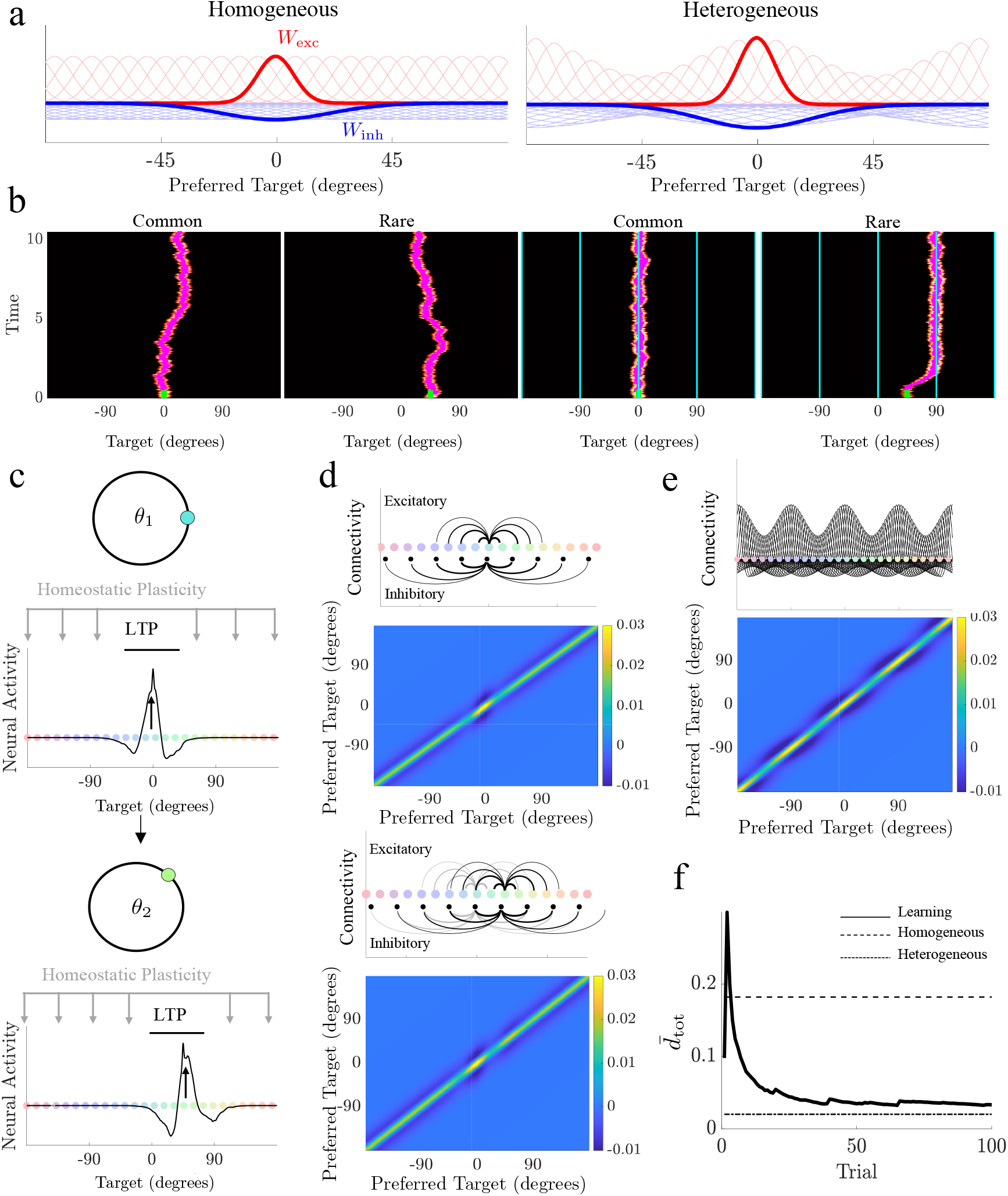
Experience-dependent modulation via long-term potentiation and homeostatic plasticity reduces distortion of encoded stimulus values. **a.** Localized excitatory (*W*_exc_) and inhibitory (*W*_inh_) synaptic weights associated with preferred item features in a neural field encoding a delayed estimate. Example of of synaptic weight originating from neuron with preference *θ* = 0 shown in bold. Heterogeneous neural networks are modulated so synaptic footprints originating from peaks of *P*_env_(*θ*) are stronger. **b.** Example bumps of sustained neural activity over 10s delay period originating at common and rare targets in a homogeneous (left) and heterogeneous (right) case. Cyan traces in heterogeneous plots denote attractor locations with enhanced synaptic weights. Stimulus input duration shown in green. **c.** Experiencedependent learning results from long-term potentiation (LTP) of neurons with a preference for the previous target value and homeostatic reduction of connectivity elsewhere. **d.** Learning breaks the symmetry of the spatially-dependent weight kernel, creating enhanced peaks originating from neurons activated across trials. **e.** After many trials, the weight matrix recovers the static heterogeneous synaptic structure. **f.** Average total distortion over time decreases as the experiencedependent neural field model learns the environmental distribution. Parameters for all sub-figures as listed in Methods Table 3.

Spatially-varying heterogeneity (learned or prescribed) acting on the synaptic connectivity breaks the symmetry of the spatial synaptic footprint emanating from each neuron. When heterogeneity is strongest at the peaks of the environmental distribution, attractors are created (Fig. **5a**; right) that bias bumps to drift toward the most common stimulus locations (Fig. **5b**; right). This relationship between the increase in synaptic efficacy and the formation of attractors can be made mathematically precise via direct asymptotic analysis (see Methods). As such, there is a direct relationship between the stochastic dynamics of a bump’s position and the particle models we have discussed already. In short, the introduction of synaptic heterogeneity effectively reshapes an energy landscape that determines the bump position. While this reduces bump wandering when encoding common stimulus values, bumps drift more when instantiated at rare targets, causing larger errors as they are drawn toward attractor locations. As with the particle models, total mean distortion of input stimulus values during the delay is reduced by spatial heterogeneity aligned to the environmental prior (Fig. **S8**).

We next identify a neuromechanistic learning rule that can modulate synaptic strength based on experience, reshaping the effective energy landscape along which the bump’s position evolves: Synapses emanating from activated neurons (those encoding the stimulus value) are potentiated [22], while homeostatic plasticity compensates for this local increase in synaptic strength by reducing synaptic strength elsewhere throughout the network (Fig. **5c**). This rule comes from ample evidence for physiological mechanisms supporting long-timescale presynaptic potentiation throughout the nervous system [44, 45] and homeostatic mechansisms that can prevent runaway positive feedback loops of excitation and potentiation [46, 47]. Such synaptic modulation leads to an increase in connectivity strength at the target location of each trial and a reduction of synaptic efficacy elsewhere (Fig. **5d**). Updates occur iteratively, so synaptic plasticity modulates weight functions across long timescales to reflect the environmental prior (Fig. **5e**). As with our particle models, once the neural network learns the environmental prior through experience, it maintains delayed estimates with reduced total mean distortion compared to homogeneous networks with fixed synaptic structure (Fig. **5f**).

## Discussion

We have demonstrated that systematic biases observed in human subjects’ delayed estimates can be attributed to environmental experience, specifically corresponding systematic variations in the frequency of stimulus feature values. Our work identifies a learning mechanism that can be implemented in reduced models and physiologically motivated neural circuit models and is validated by human response data. This moves beyond prior work which primarily proposed analogous attractor-based models with fixed energy landscapes [16, 29]. Our analysis lends credence to the claim that errors reflect a gradual inference of the environmental stimulus prior.

Beginning with a simplified model of delayed-estimate degradation, we confirm that systematic response biases can be induced by breaking the symmetry of the energy landscape that shapes the evolution of the delayed estimate over time. Such symmetry-breaking stabilizes memories at attractor locations that, when aligned to peaks in the environmental prior, reduce response error at common stimulus values at the expense of larger errors for rare feature values. Overall, total mean distortion of responses is reduced in models with aligned heterogeneous energy landscapes, compared to those with flat landscapes, given the higher propensity of common input stimulus values. Experience-dependent learning of the environment can be implemented neuromechanistically via long-term potentiation that enhances recurrent excitation in neurons encoding common stimulus values and homeostatic plasticity that regulates connectivity across the neural population. Responses from human subjects are better matched to models that learn environmental priors than those that do not, particularly if the task environment does not well match their baseline beliefs. Thus, subjects confronted with environments that deviate from their priors appear to dynamically update their beliefs based on experience, supporting our hypothesis that systematic biases are learned via experience-dependent plasticity.

Our work supports previous findings that response variability can be reduced in neuronal networks with spatial heterogeneity in synaptic connectivity, even if the stimulus probability is uniformly distributed across values [16,28–30], and extends these findings to measure the efficiency of such codes when stimulus priors are non-uniform. While others have shown repulsive effects in persistent activity codes when implementing heterogeneous priors and computing efficient codes [6,48], our models suggest synaptic heterogeneities should be aligned with peaks in the environmental prior to incorporate stronger connectivity at common stimulus values. These differences in results can be explained by applying two different underlying mechanisms to learning the environmental prior: “anti-Bayesian” biases can emerge from redistributing the frequency of neural tuning preferences such that more neurons have stimulus preferences near common feature values, and the stimulus estimate is biased away from common values due to the higher variance in the estimation of rare values [48]. In contrast, our work suggests that synaptic plasticity modifies connectivity to encode environmental priors. In humans, results on systematic working memory biases are mixed [21,49–51], suggesting that subjects do not necessarily use consistent or optimized strategies. Our work supports this finding by identifying a number of different models that best match individual subject’s responses, many of which produce suboptimal results.

For example, subjects’ use of the distractor item as part of their updating procedure suggests that experience-dependent updates could occur during stimulus observation, rather than after subjects’ response as suggested by work on short-term serial biases [8,22]. Representations of memoranda in multiple item working memory tasks have also been shown to interact, sometimes causing additional errors in memory [18,52,53] or reducing cardinal biases [54]. Notably, multi-items were presented sequentially in [54], implying that both short-term plasticity rules [22] and multi-item interactions, such as swapping errors [55], may work in conjunction to produce suboptimal strategies. Future work may consider how multi-item working memory tasks impact experience-dependent learning of task environments.

We had hypothesized that our fixed heterogeneous models would better represent subject responses when environmental priors were more aligned to the population biases (offsets closer to the population bias peaks), because these environments would require less updating to subjects’ environmental beliefs. In contrast, the population of subjects whose strategies are best described by fixed heterogeneous models have a wide range of assigned offsets, while most subjects described by learning models have assigned offsets that deviated from the original population biases. It is unclear whether subjects matched to static models were resistant to learning or were not given sufficient experience (i.e., trials) to adapt. Future studies could investigate the rate of learning when subjects are presented with stimuli drawn from heterogeneous environmental priors to identify the causes for subject-model variability.

Our work has established and validated a novel mechanistic hypothesis to describe how people infer the distribution of environmental stimuli and its impacts on their delayed estimates. Our results support recent findings on training-induced changes in prefrontal cortex [56], suggesting learning over longer timescales can have substantial stimulus-specific impacts in working memory. Moreover, our work posits that limitations and biases in working memory are not necessarily suboptimal, but can be motivated by efficient coding principles and modulated by environmental inference processes. These findings establish a correspondence between environmental inference and working memory that reveals a deeper understanding on the role of working memory in cognitive processes.

## Materials and Methods

### Particle Model

We described the models here in radian coordinates (i.e., the distance around the ring is 2*π* rather than 360 degrees), but all figures were plotted by rescaling to degrees deg = (180/*π*) · rad. All particle models with fixed energy landscapes used

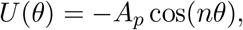

in which *A_p_* described the amplitude and *n* described the number of wells (attractors). The homogeneous model was recovered when taking *A_p_* = 0. Particle movement was simulated using a stochastic differential equation

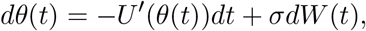

which incorporated noise as a Wiener process with increment *dW*(*t*). Numerical simulations were performed using the Euler-Maruyama scheme in which the values for *θ* were discretized to 1 degree (*π*/180 radian) bins and time was discretized to 10ms bins. To recover the effects of drift alone –*U*′(*θ*(*t*))*dt*, we set *σ* = 0. All parameter values listed in Table 1 were used unless otherwise stated.

**Table 1:** Parameter values for particle models.

| Variable | Value |
| --- | --- |
| $\sigma$ | 0.05 |
| $A_p$ | 1 |
| $n$ | 4 |
| $T_{\text{Delay}}$ | 5 |
| $\beta$ | 8 |
| $h$ | 0.25 |
| $s$ | 5 |

### Distortion

The mean distortion for a given input stimulus value was computed as

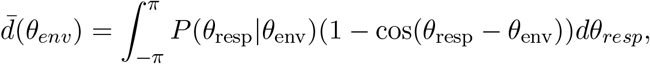

using Monte Carlo sampling. To compute stimulus-specific distortion for a given particle model and environment, *θ* was binned and simulations were used to compute 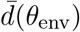 for the bin (*N*_sim_ = 10^5^ per bin). Bootstrapping procedures were used to resample distortion and compute the standard deviations (*N*_boat_ = 1*e*3). To compute the the effects of distortion purely based on diffusion for heterogeneous models, simulations were sampled at a single attractor/common stimulus value (*θ* = 0), while drift-only dynamics were simulated with *σ* = 0.

Total mean distortion across all stimulus values in an environmental prior was computed

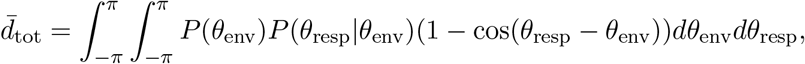

which can be approximated by Monte Carlo simulations with initial conditions sampled from the environmental distribution *P*_env_ (*N*_sim_ = 10^5^).

### Conditional and Marginal Distributions

The conditional probability *P*(*θ*_resp_|*θ*_env_) was computed by simulating the distribution of responses (particle end locations) for each discretized *θ*_env_ value (*N*_sim_ = 10^5^). Marginal distributions of the response *P*(*θ*_resp_) were computed by averaging the discrete conditional probability solutions relative to the known environmental distribution *P*(*θ*_env_).

### Relating the energy landscape to an experience-based posterior

An experience-based posterior can be related to the stationary distribution of a particle on an energy landscape associated with Eq. (2). The equivalent Fokker-Planck equation describing the evolution of the distribution *p*(*θ, t*) of possible particle positions *θ* at time *t* assuming a potential function *U*(*θ*) was

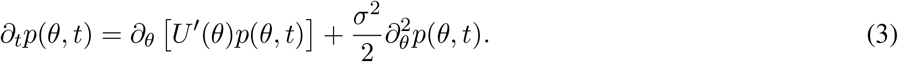

We derived the form of *U*(*θ*) that led to a stationary density that corresponds to a particular posterior *L*(*θ*) in the limit *t* → ∞ in Eq. (3). The stationary density 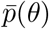 was analogous to a posterior *L*(*θ*) since it is the probability density that the system represents when there is no information about the current trial’s target remaining. Thus, we derived an association between 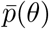 and *U*(*θ*) to identify how the energy landscape function *U*(*θ*) should be tuned so that 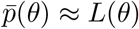. In the limit *t* → ∞, we found Eq. (3) becomes

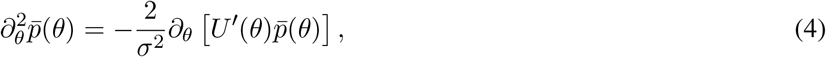

a second order ordinary differential equation with solution

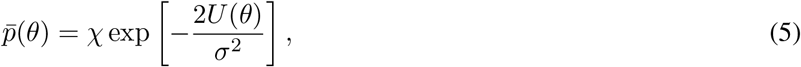

where *χ* is a normalization factor. Thus, to match 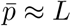, we need

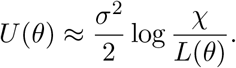

We assumed that we were in the limit of weak heterogeneities, so the deviation of the function *L*(*θ*) from flat will be weak, and 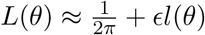 (where 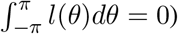, which allowed us to make a linear approximation

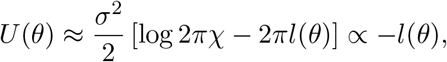

Thus, we removed the constant shift and were only concerned about the proportionality of the energy landscape to the negative of the variation *l*(*θ*) in the posterior.

### Experience-dependent particle model

To incorporate learning into the particle model, we updated its energy landscape based on the history of experienced stimulus values according to the equation

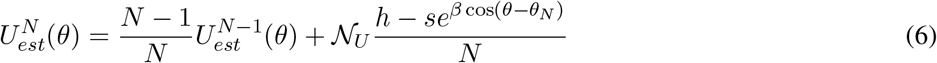

which incorporated a von Mises distribution centered at the location of the true stimulus value *θ_N_* on trial *N* with *s* as the scaling factor, *h* the shift, and *β* the spread. This energy landscape update was meant to represent the trial-by-trial probabilistic update to the stimulus distribution estimate. The mean distortion and the particle landscape were updated iteratively on each trial, such that for each stimulus, the distortion was computed and included in the running average. All parameter values listed in Table 1 were used unless otherwise stated.

The additive update of the particle landscape linearly approximated the typical multiplicative scaling of posterior updating based on successive independent observations. To demonstrate how the updating rule for the energy landscape is related to Bayesian sequential updating of the posterior, recall that that enforcing *U*(*θ*) ∝ –*l*(*θ*) ensured an energy landscape aligned with the learned posterior. Thus, we derived an approximate inferred distribution of possible future stimulus target values, updated based on the observed history *θ*_1:*N*_. We assumed that when an observer sees a target value *θ_N_*, they inferred that subsequently similar values are more likely, according to the von Mises distribution

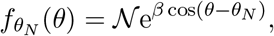

where 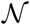 was a normalization factor. We noted this was self-conjugate (*f_θ′_*(*θ*) ≡ *f_θ_*(*θ*′)). We will also assumed that 0 < *β* ≪ 1, so the variation in *f_θ′_*(*θ*) was weak, which allowed us to approximate 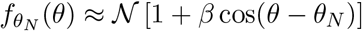. Sequential analysis then could determine how a posterior for future observations should be updated based on each observed target. Take *p_N_*(*θ*) = *p*(*θ*|*θ*_1:*N*_) to be the posterior based on past observations *θ*_1:*N*_ which can be computed directly as the product of probabilities

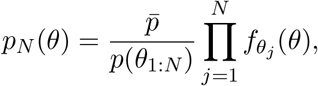

where 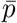 is the uniform distribution and we have utilized the self-conjugacy of *f_θ′_*(*θ*) ≡ *f_θ′_*(*θ′*). We used the linearization of the likelihood function and truncated to linear order in *β* to find

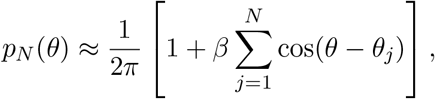

which, with the approximate formula for *f_θ_N__*(*θ*), can be written as

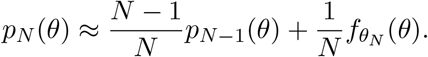

Lastly, noting the proportional relationship of the desired energy landscape to the posterior, *U_N_*(*θ*) ∝ –*l_N_*(*θ*), we found that the appropriate update for the energy landscape to match this iterative additive update of the posterior was

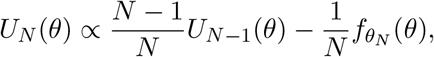

which we could rewrite using the full form of *f_θ_N__*(*θ*) plus a shift to obtain Eq. (6).

Thus, in the long-term limit (as *N* → ∞), the energy landscape convolved the environmental prior *P*_env_(*θ*) against the negative of the likelihood function:

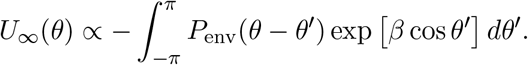

Given that the environmental prior had the form 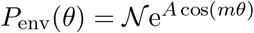, we then made the approximation 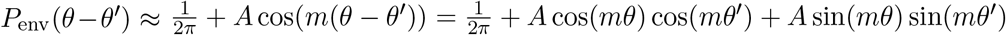, so we could compute

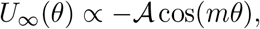

where 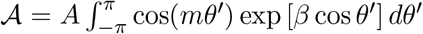 and the other term vanished due to its odd symmetry. This was consistent with the form of the fixed heterogeneity we used to align with this environmental prior.

### Human data

Response data from a delayed estimation task was taken from [16], experiment 2, with permission. The task was administered to 120 consenting subjects with normal color vision in Amazon Mechanical Turk who performed and achieved minimal engagement. Each trial within the task presented a subject with two colored squares simultaneously for 200ms after which time they disappeared and a delay of 500 ms (100 short trials) or 4000 ms (100 long trials) ensued prior to a response being cued by presenting an outlined square in the location of one of the two previous prompt (implicit identification of the target object). Participants then provided an estimate of the cued color by using a mouse to drag a small circle around a ring of colored continuum. Each item had a 50% chance of being drawn from the biased distribution. The biased distribution included 4 peaks spanning 20 degrees, equally spaced about the circle. The offset of the stimulus peaks were picked uniformly and randomly and assigned independently to each subject. The location of the population bias was identified based on the peaks in response frequency across the population of human subjects observed in experiment 1 from [16], which probed subjects to report a color drawn from a uniform distribution but subject showed preferences in the reports.

### Subject model fitting

We fit subject responses to 8 different particle models and identified the most likely model using cross-validation:

1. **Flat potential** (1 free parameter) in which the particle dynamics were only influenced by diffusion

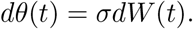
2. **Static Heterogeneous** (3 free parameters) in which the particle was subject to drift and diffusion, parameterized by the *A_p_* (amplitude), *n* (number of wells), and noise *σ*,

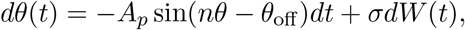

where *θ*_off_ was the offset assigned to a subject by the experiment (not fit).
3. **Offset Heterogeneous** (4 free parameters) included all of the above parameters but incorporated a free parameter for the offset value 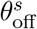, such that a subject could use a model not aligned to their assigned offset *θ*_off_.
4. **Dual Heterogeneous** (5 free parameters) assumed that subject response were governed by an energy landscape determined by two frequencies (*n*_1_ and *n*_2_) with amplitudes *A*_1_ and *A*_2_, and assuming the offset to be at the assigned location. The stochastic dynamics of the particle were described

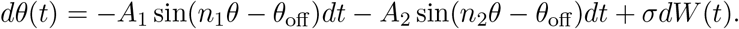
5-6. **Target-only Learning** (3 free parameters) assumed the energy landscape was updated on each trial as described in the experience-dependent particle model section above, by adding an inverted von Mises distribution centered at the target location with free parameters for *β* and *s*. Noise was parameterized by *σ* as before. The initial landscape was either chosen to be (a) flat *U*_0_(*θ*) = 0, or (b) heterogeneous *U*_0_(*θ*) = –*A_p_* cos(4(*θ* – *θ*_off_)) with *A_p_* = 1 and *θ*_off_ aligned to the offset of the established population biases described above.
7-8. **Target + Distractor Learning** (3 parameters) models were implemented equivalently to the “Target-only” model but were updated by adding two inverted von Mises distributions to the energy landscape on each trial, one centered at the target and the other centered at the distractor stimulus value.

Models were fit to each subject’s set of responses using 5-fold cross-validation performed for short and long delay trials separately. For each subject-model, we tested 100 parameter sets, selected uniformly from a bounded domain for each parameter (Table 2), on 80% of the trials, running 100 simulations with each set of parameters for each trial. The mean squared error (MSE) between each simulated and subject response were computed, and the parameter set with the lowest MSE was selected for that subject-model pair. We then simulated responses for the final 20% of trials using the selected parameter set and computed the MSE for these trials. This process was performed 5 times, testing all trials. The testing-set MSEs were then averaged, and the model with lowest mean testing-set MSE was selected for each subject.

**Table 2:** Parameter ranges for human response model fitting.

| Model Class | Variable | Bounded Domain |
| --- | --- | --- |
| Flat, Fixed, Learn | $\sigma$ | 0.01 – 0.2 |
| Fixed | $A_p/A_1/A_2$ | 0.1 – 2 |
| Fixed | $n/n_1/n_2$ | 1 – 12 |
| Fixed | $\theta_{\text{off}}^s$ | $0 - \pi/2$ |
| Learn | $\beta$ | 1-10 |
| Learn | $s$ | 1-10 |

### Neural Field Model

We used a lateral inhibitory neural field model on the ring [29,43,57], in which the locations of neurons corresponded to their preferred stimulus value

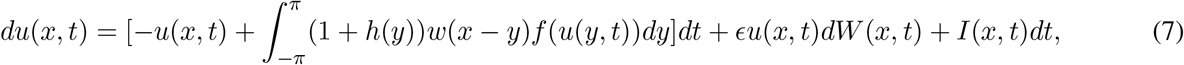

where *u*(*x, t*) was the mean activity at the location *x* ∈ [–*π*, *π*]. Spatial connectivity was described by

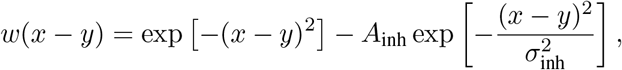

combining both local excitation and broad inhibition, where *A*_inh_ was the strength of inhibition and 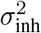 described the inhibitory spread. Heterogeneity in connectivity was described by the function *h*(*y*), which could be learned or fixed. Fixed periodic heterogeneity was incorporated by taking *h*(*y*) = *A_n_* · cos(*ny*), where we bounded *A_n_* ∈ (–1,1) to > preserve excitatory/inhibitory polarity.

The firing rate nonlinearity *f*(*u*(*y, t*)) was taken to be a Heaviside function

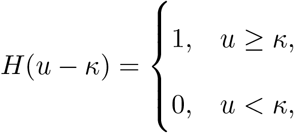

in which *κ* described the firing rate threshold.

Noise *ϵu*(*x, t*)*dW*(*x, t*) was weak, multiplicative, and driven by a spatially-dependent, white-in-time, Wiener process with the spatial filter that decayed with distance |*x* – *y*|:

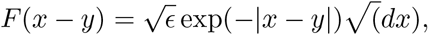

and *ϵ* described the noise strength.

Input to the network corresponding to the true location of the stimulus target at location *x*_targ_ was given by

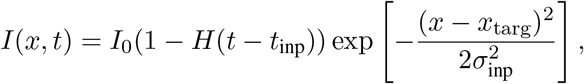

where *I*_0_ was the strength of the input and 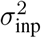 parameterized the width of the input. Note, the location *x*_targ_ was sampled from the environmental distribution *P*(*x*_env_) as described above to comprise a long sequence *x*_1:*N*_ across trials.

Neural activity evolved by applying Euler-Maruyama iterations to the timestep *dt* and Riemann integration with dx to the integral in the discretized version of Eq. (7). The bump’s centroid was then identified as the peak in neural activity at each time *θ*_cent_(*t*) = argmax_*x*∈[–π,π]_*u*(*x, t*). All model parameters are given in Table 3 and were selected to ensure bumps would not extinguish prior to the end of the delay period. Responses for each trial were reported as the location of centroid at the end of the delay period *θ*_cent_(*T*).

**Table 3:** Neural Field Parameter Values.

| Variable | Value |
| --- | --- |
| $A_{\text{inh}}$ | 0.35 |
| $\sigma_{\text{inh}}$ | 3 |
| $A_n$ | 0.4 |
| $n$ | 4 |
| $\kappa$ | 0.1 |
| $\epsilon$ | 0.5 |
| $I_0$ | 1 |
| $t_{\text{inp}}$ | 0.5 |
| $\sigma_{\text{inp}}$ | 1 |
| $T_{\text{delay}}$ | 10 |
| $A_{\text{inp}}$ | 1 |
| $s_{\text{learn}}$ | 0.1 |
| $dt$ | 0.1 |
| $dx$ | 0.036 |

### Linking the neural field and particle models

The dynamics of bump solutions to Eq. (7) can be reduced to first order to describe how their position 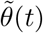 evolved over time, roughly approximating the centroid (peak location of neural activity). A reduced stochastic differential equation can be derived describing how this position evolves in time due to noise, inputs, and heterogeneity in the weight function. Technical details for such calculation can be found in [22,38]. Here we give a brief sketch of such analysis, to demonstrate the tight mathematical link between our particle models and the stochastic dynamics of bump solutions to our neural field equations.

Ignoring noise (*ϵ* → 0), heterogeneity (*h*(*y*) → 0), and input *I* → 0, Eq. (7) had bump solutions 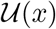 that satisfied the equation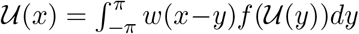 [38]. This bump was marginally stable and lay on a continuous attractor, so it could be placed at any position [–*π, π*] [43]. Without loss of generality, we assumed this position was initially *x* = 0, we could track dynamics of the bump’s position 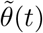 once noise, heterogeneity, and input were reintroduced by deriving a hierarchy of equations for the expansion 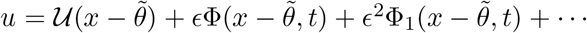. Enforcing solvability of this hierarchy introduced a condition requiring the sum of the noise, input, and heterogeneity to be orthogonal to the nullspace *φ*(*x*) of the adjoint of the operator that comes from linearizing Eq. (7) about the bump solution. The result was a drift-diffusion equation whose drift was determined by the energy landscape invoked by both the synaptic weight heterogeneity and input

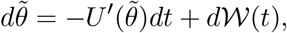

precisely the form of Eq. (2), where the drift had contributions from the weight heterogeneity and input

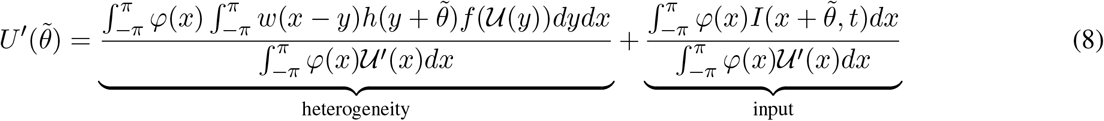

and the Wiener process noise 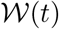 had zero mean and variance

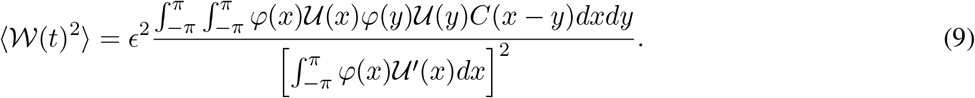

The heterogeneity and input introduced an energy landscape that steers the position 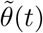 of the bump as it responds to noise fluctuations. As shown in [22,38], by dropping the input term and considering a Heaviside nonlinearity *f*(*u*) = *H*(*u* – *κ*), 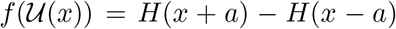 and *φ*(*x*) = *δ*(*x* – *a*) – *δ*(*x* + *a*) where *a* was the half-width of the bump such that 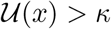 for *x* ∈ [–*a, a*] and 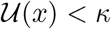 otherwise and *δ* was a Dirac delta function. As such, we could simplify the energy landscape gradient formula to find

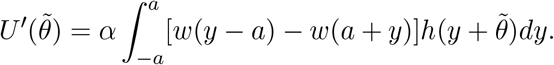

#### Approximation with Fourier modes

Note that by decomposing the even weight function into its Fourier series, we have

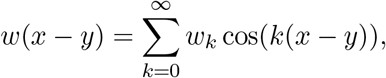

which allowed us to write

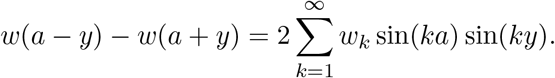

In a similar way, we could decompose the function describing the heterogeneity in the weight

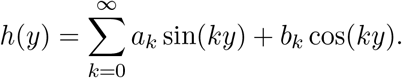

Approximating by the dominant Fourier mode (assume it is even, *m* = argmax*_k_b_k_*), we took *h*(*y*) ≈ *b_m_* cos(*my*). Integrating against the difference of the shifted homogeneous weight function, then we found 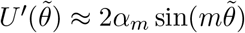 and thus 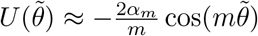, where

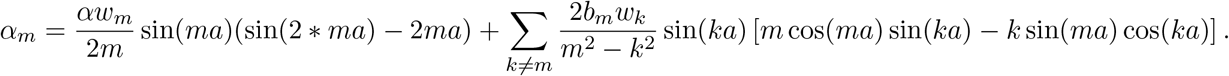

Note also that as *m* and *k* differ more, the coefficient in the sum will decrease, suggesting the dominant terms from the series description of *w* will be those for the modes *k* indexed close to *m*. Thus, a scaling of the dominant Fourier mode of the weight heterogeneity well approximated the energy landscape associated with the bump’s stochastic motion.

#### Narrow bump approximation

Assuming the bump width was narrow compared to the length scale of the heterogeneity, we could estimate the integral using the trapezoidal rule

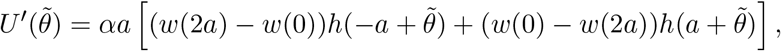

so by expanding the even weight function *w*(2*a*) ≈ *w*(0)+2*a*^2^*w*″(0) as well as linearizing the heterogeneity 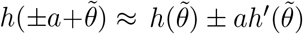, we obtained

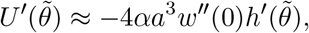

and thus

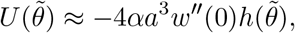

so the energy landscape generated for the bump position from weight heterogeneity *h*(*y*) was approximately proportional to the negative shape of the heterogeneity. As such, any neurons whose emanating synapses were potentiated/depressed then attract/repulse the bump.

### Plasticity rules in neural field model

To include experience-dependent learning into our neural field model, we allowed the function that modulated the synapses emanating from each neuron to evolve (1 + *s*(*y,t*)), updating with each trial *N* based on presynaptic neural activation 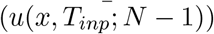 while the network received input (i.e., at the stimulus location in trial *N* – 1, *x*_*N*–1_),

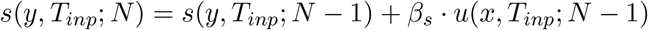

where *β_s_* was a scaling factor for long-term plasticity. Such a rule implemented an activity-dependent form of pre-synaptic potentiation, deemed transmitter-induced long-term plasticity by [46] and for which multiple mechanisms have been proposed [47,58,59]. Equivalently, this is a slow form of short-term synaptic facilitation, emerging in neural field models the same way we have included it here [22, 37]. Inclusion of homeostatic mechanisms was based on ample evidence showing it capable of preventing runaway potentiation [46,60]. In particular, we considered a mechanism that provides a saturation threshold setting an upper limit on potentiation [61]. Since potentiation would otherwise drive network strength above this threshold, mathematically this amounted to always normalizing peak synaptic strength via the computation

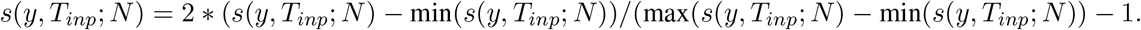

As in the case of the energy landscape, we could determine the long-term limiting heterogeneity *s*_∞_(*y*) resulting from the learning rule combined with an environmental prior *P*_env_(*θ*). Approximating the shape of the instantiated bump by a von Mises distribution centered at the location of the stimulus value on each trial and assuming weak modifications to the heterogeneity, the long time limit gave

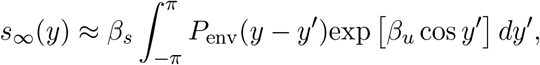

and, by making the approximation 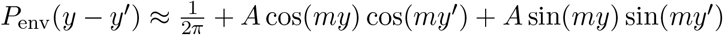, then

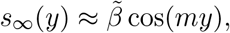

consistent with the expected form of synaptic heterogeneity and resulting energy landscape.

## Funding

We thank Matthew Panichello and Krešimir Josic for their valuable feedback on the manuscript. We thank Matthew Panichello and Timothy Buschman for supplying the previously published human data analyzed here. This work is funded by National Institutes of Health grant 1R01EB029847-01.

## Author Contributions

Conceptualization: TLE, ZPK

Methodology: TLE, ZPK

Investigation: TLE

Visualization: TLE

Supervision: ZPK

Writing-original draft: TLE

Writing-review and editing: TLE, ZPK

## Competing Interests

The authors declare that they have no competing interests.

## Data and Materials Availability

The data used to generate the figures and code developed for all proposed models has been made available on github.com/teissa/HeterogeneousWorkingMemory.

## Supplementary Materials

**Figure S1:**
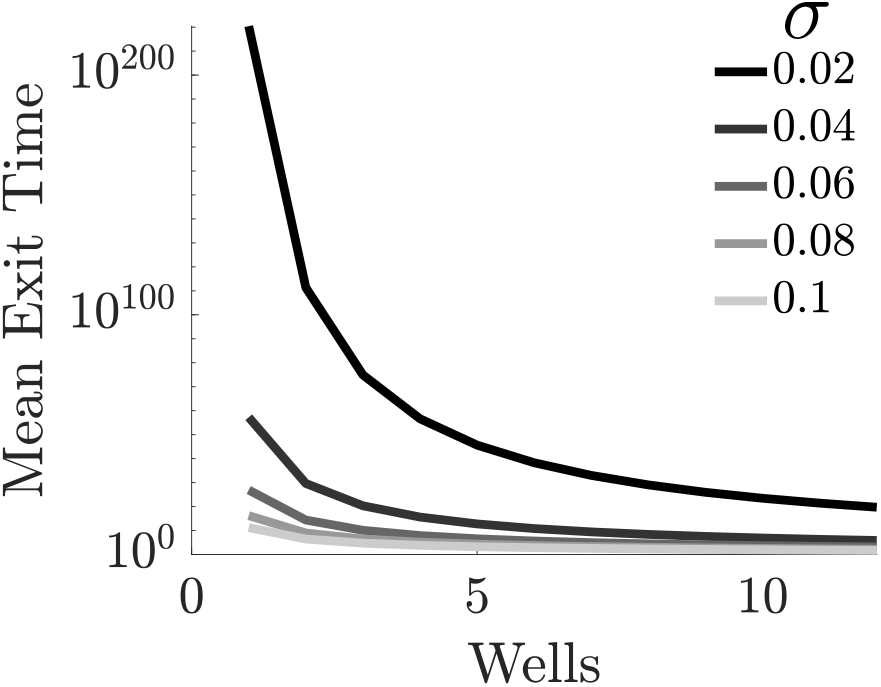
Mean exit time for a particle to leave the current well of attraction. Computed as in [29, 42]. Parameters as listed in Methods Table 1.

**Figure S2:**
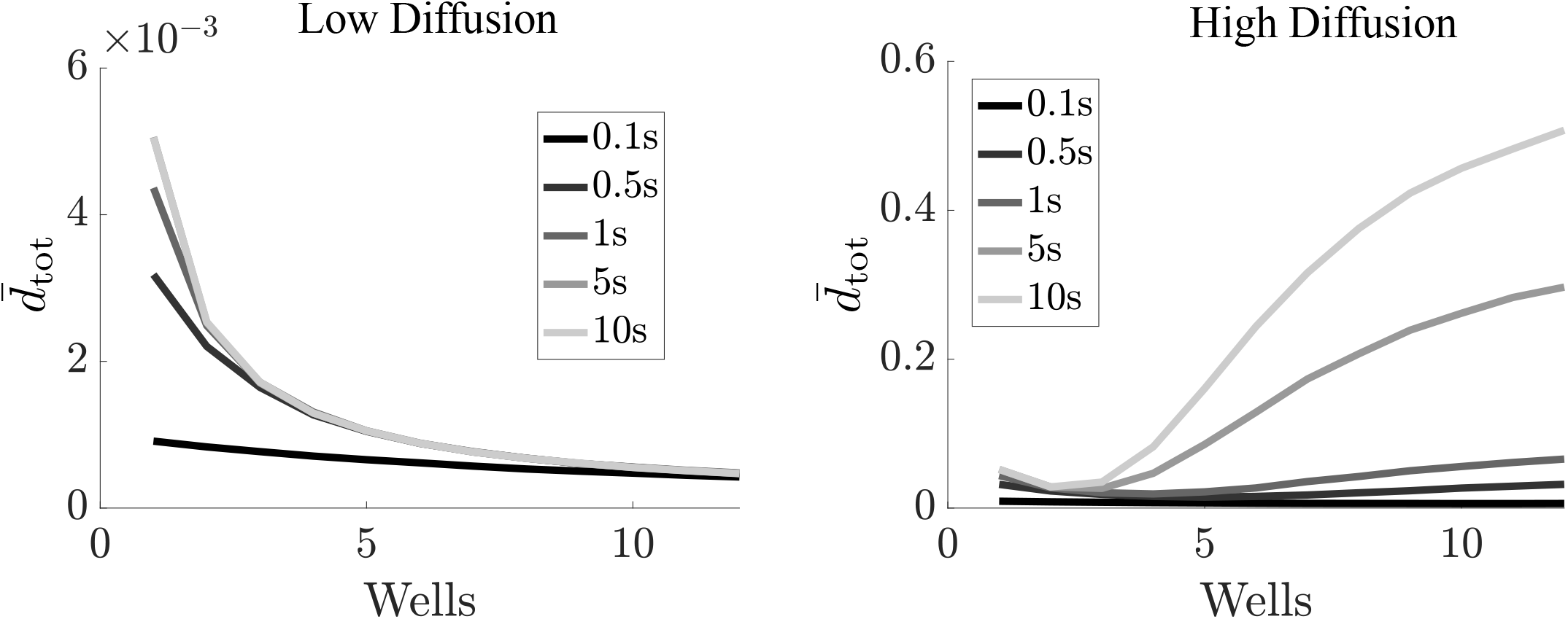
Total mean distortion in a heterogeneous landscape with only diffusion (targets sampled at attractor points), low diffusion (*σ* = 0.01, left) and high diffusion (*σ* = 0.1, right). Parameters as listed in Methods Table 1.

**Figure S3:**
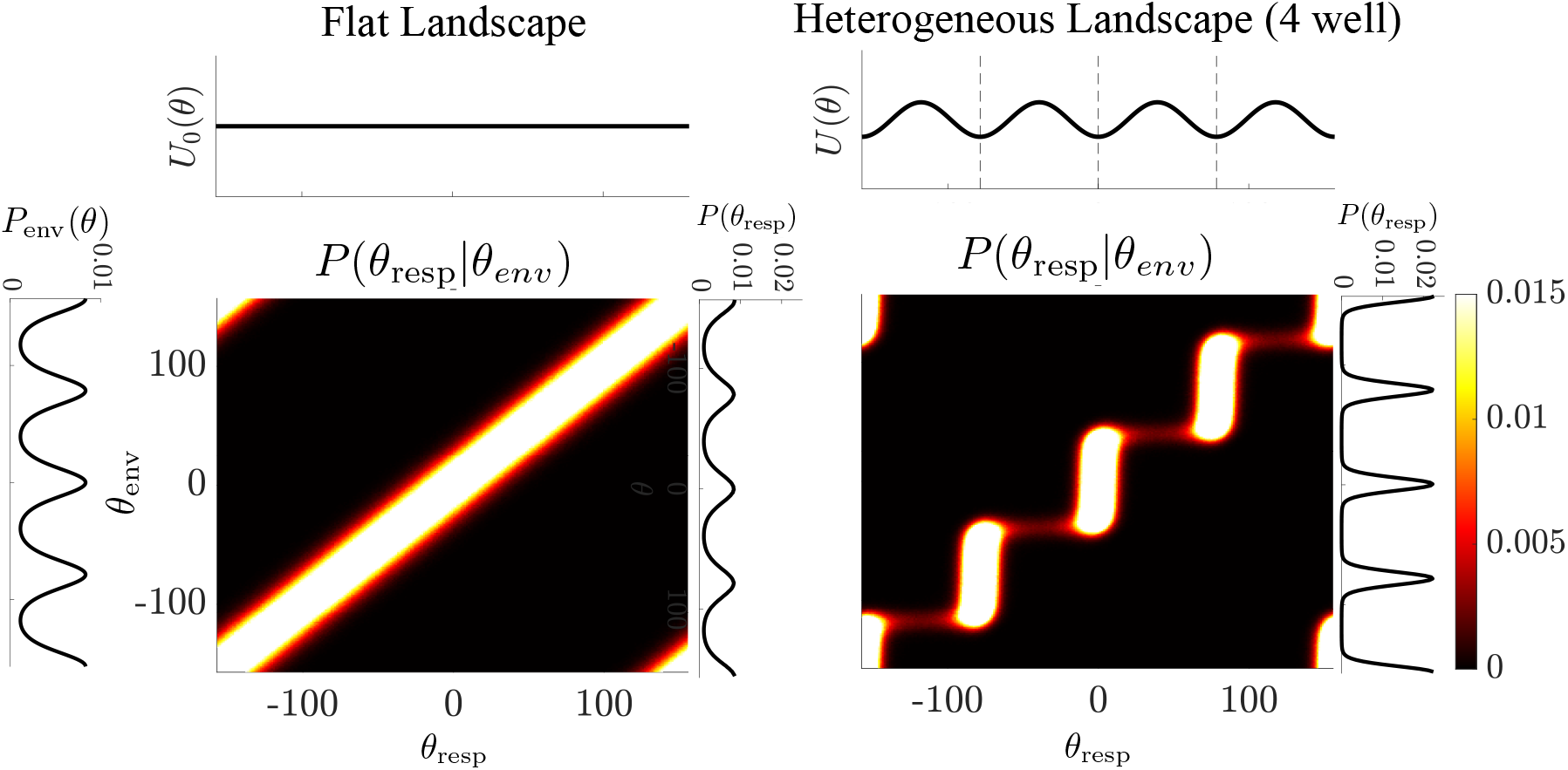
Comparing energy landscapes (*U*(*θ*)) and heterogeneous feature value distribution (*P*_env_(*θ*)), we find the conditional probability of response *P*(*θ*_resp_|*θ*_env_) and the marginal probability of response *P*(*θ*_resp_) for particle models with homogeneous and heterogeneous (four wells at environmental distribution peaks) landscapes. Parameters as listed in Methods Table 1.

**Figure S4:**
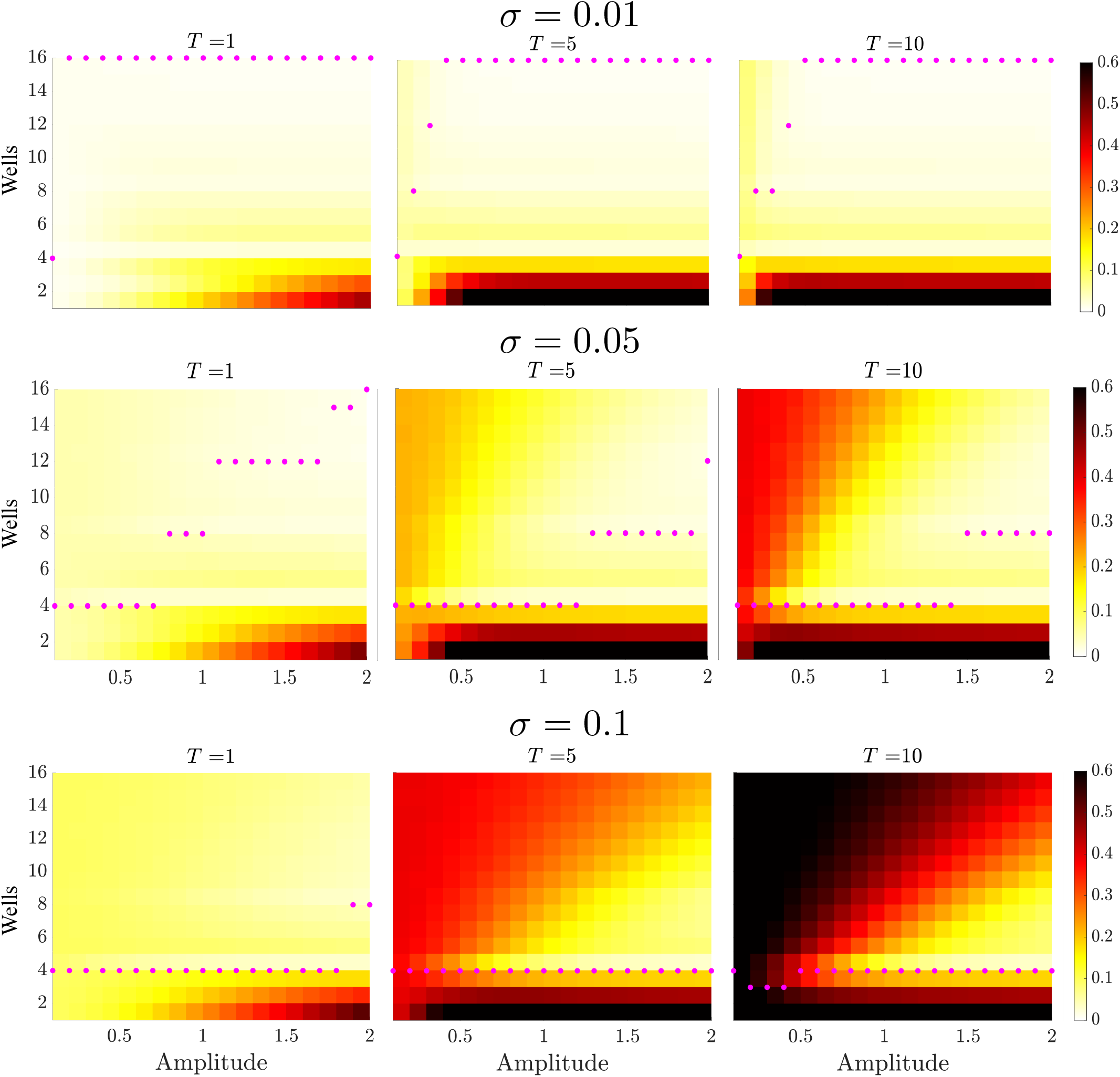
Total mean distortion as amplitude and number of wells was varied for three example delay periods. Optimal particle model identified based on minimum mean distortion (magenta dots). Top: Low diffusion (*σ* = 0.01) leads to a optimal models with higher number of wells. Center: Moderate diffusion (*σ* = 0.05) leads to optimal models with a variable number of wells based on amplitude, often harmonics of the number of environmental peaks. Bottom: High diffusion (*σ* = 0.1) leads to a optimal models with lower number of wells.

**Figure S5:**
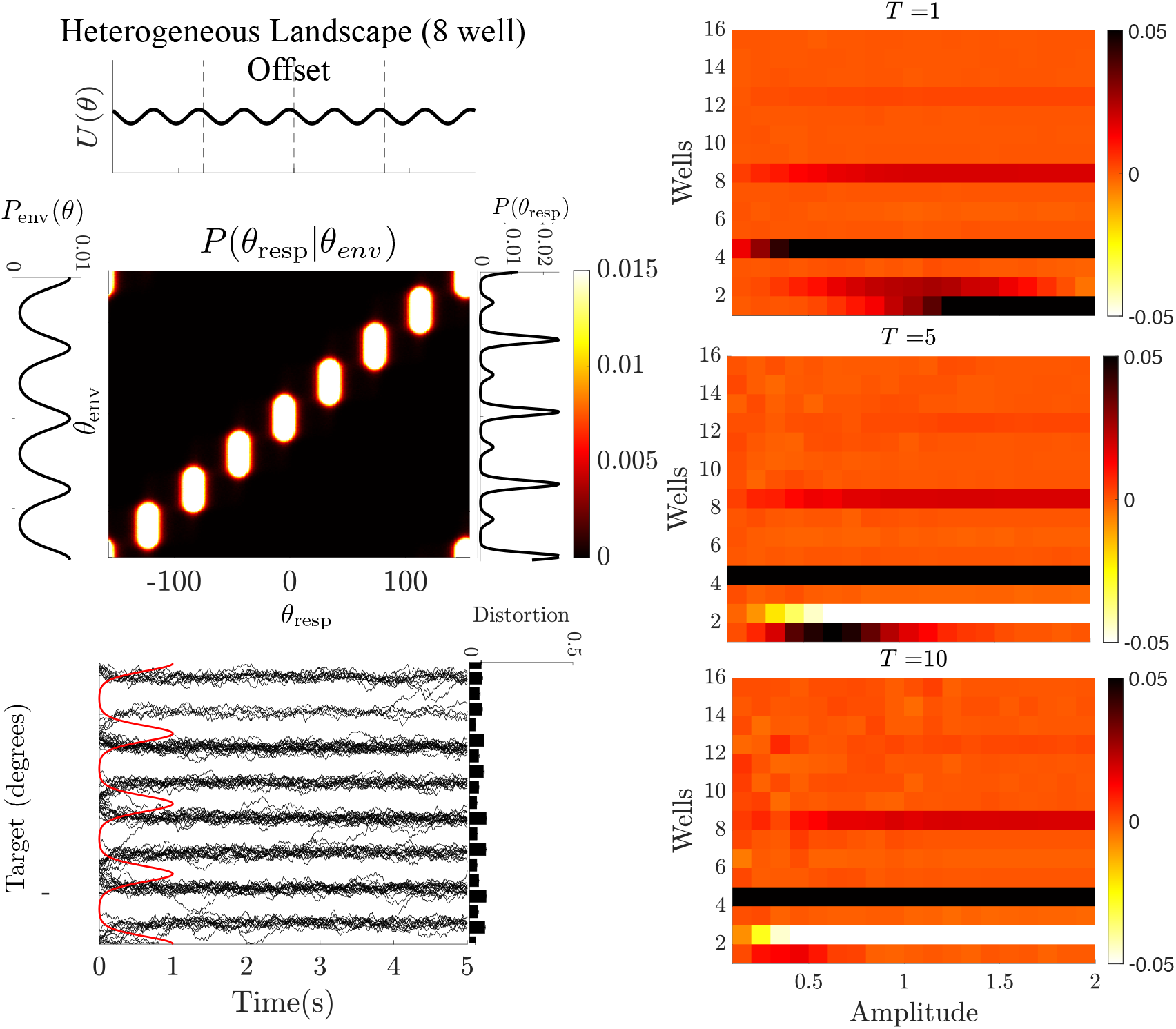
Left: The conditional and marginal probabilities when the heterogeneous particle model has more wells than the environment and offset from the peak locations. This offset leads memoranda to drift to offset locations and shows moderate distortion for all values of *θ*. Right: Total mean distortion in offset heterogeneous particle models as compared to non-offset models for moderate diffusion (*σ* = 0.05). Positive values corresponds with higher levels of distortion in offset models. Parameters: *T*_Delay_ = 1, *n* = 8 offset = 45, all others as listed in Methods Table 1.

**Figure S6:**
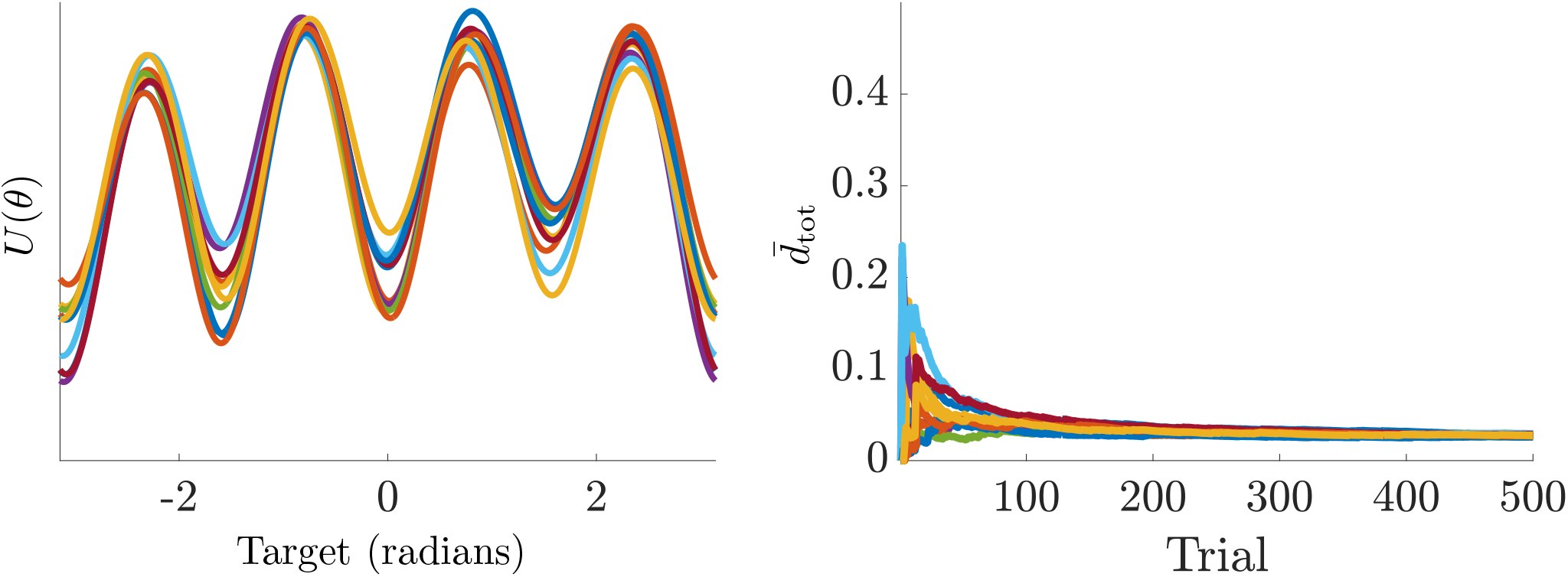
Ordering of observations in the learning particle model does not change the shape of the learned landscape or overall distortion. Left: 10 iterations of the learning model with the same observations but randomized permutations produce potential landscapes with the same shape but differing amplitudes. Right: 10 iterations of the learning model with no diffusion (drift only) and the same observations but randomized permutations show the same overall mean distortion after many trials with minor variations in the learning rate. Parameters used: *σ* = 0, all others as listed in Methods Table 1.

**Figure S7:**
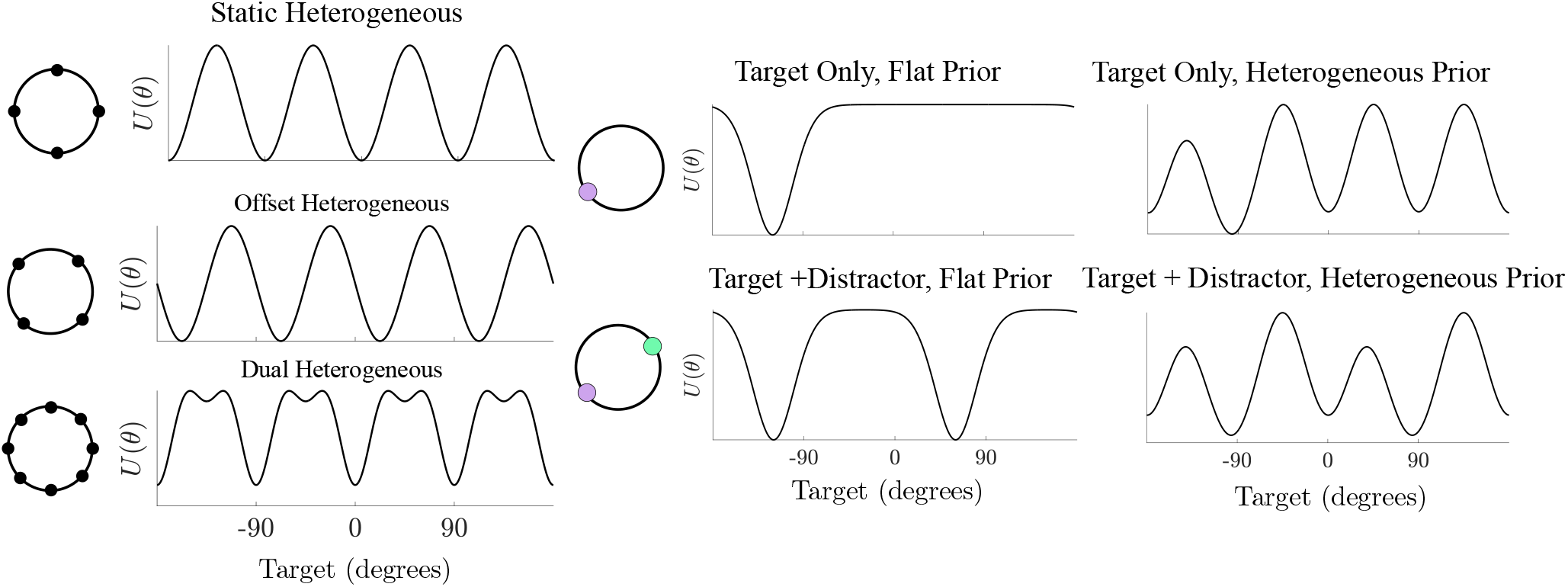
Left: All fixed heterogeneous models. Static Heterogeneous model includes three free parameters: amplitude, number of wells, and diffusion. Offset Heterogeneous includes amplitude and number of wells, diffusion, and one additional parameter for offset. Dual heterogeneous considers five parameters: amplitude and number of wells for the first component, amplitude and number of wells for the second component, and diffusion. Right: Learning particle models. Each updates iteratively based on three parameters: width of the bump, depth of the bump, and diffusion. Target-Only learning incorporated only the target prompted for response, and Target+ Distractor incorporated both items. Priors refer to initial landscape, beginning either with a homogeneous (flat) landscape or a heterogeneous landscape that matched the human population biases. Parameter ranges as listed in Methods Table 2

**Figure S8:**
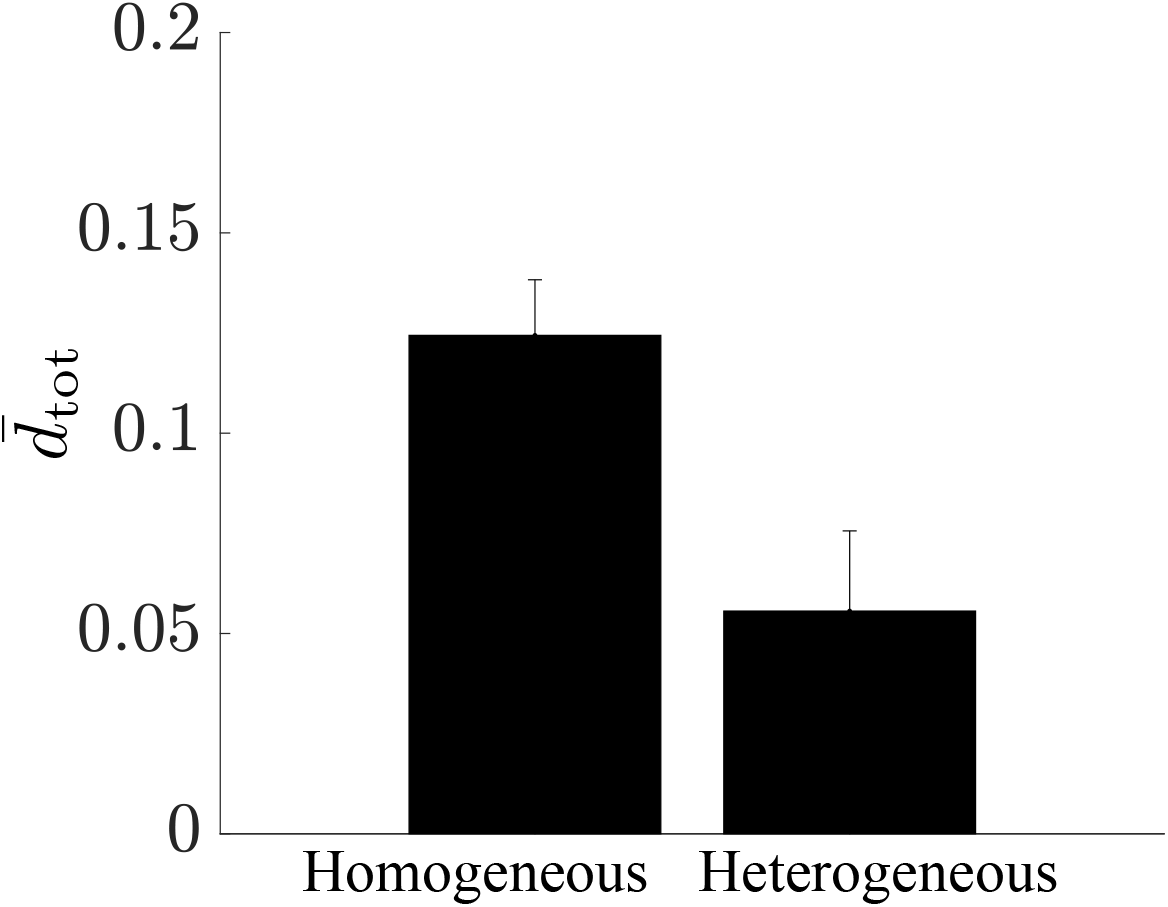
Total mean distortion in the homogeneous and fixed environment-matched heterogeneous neural field models. Bootstrapped averages (*N_Boot_* = 1*e*3) show a significant decrease in distortion for the heterogeneous synaptic connectivity. All model parameters as listed in Methods Table 3.

## Notes

https://github.com/teissa/HeterogeneousWorkingMemory

### Competing Interest Statement

The authors have declared no competing interest.

### Summary of Updates

This version has refined the text and updated the figures to better reflect the motivations of the study.

